# Efficient Sequence Embedding For SARS-CoV-2 Variants Classification

**DOI:** 10.1101/2023.08.24.554650

**Authors:** Sarwan Ali, Usama Sardar, Imdad Ullah Khan, Murray Patterson

## Abstract

Kernel-based methods, such as Support Vector Machines (SVM), have demonstrated their utility in various machine learning (ML) tasks, including sequence classification. However, these methods face two primary challenges:(i) the computational complexity associated with kernel computation, which involves an exponential time requirement for dot product calculation, and (ii) the scalability issue of storing the large *n × n* matrix in memory when the number of data points(n) becomes too large. Although approximate methods can address the computational complexity problem, scalability remains a concern for conventional kernel methods. This paper presents a novel and efficient embedding method that overcomes both the computational and scalability challenges inherent in kernel methods. To address the computational challenge, our approach involves extracting the *k*-mers/nGrams (consecutive character substrings) from a given biological sequence, computing a sketch of the sequence, and performing dot product calculations using the sketch. By avoiding the need to compute the entire spectrum (frequency count) and operating with low-dimensional vectors (sketches) for sequences instead of the memory-intensive *n × n* matrix or full-length spectrum, our method can be readily scaled to handle a large number of sequences, effectively resolving the scalability problem. Furthermore, conventional kernel methods often rely on limited algorithms (e.g., kernel SVM) for underlying ML tasks. In contrast, our proposed fast and alignment-free spectrum method can serve as input for various distance-based (e.g., *k*-nearest neighbors) and non-distance-based (e.g., decision tree) ML methods used in classification and clustering tasks. We achieve superior prediction for coronavirus spike/Peplomer using our method on real biological sequences excluding full genomes. Moreover, our proposed method outperforms several state-of-the-art embedding and kernel methods in terms of both predictive performance and computational runtime.

## 1 Introduction

The global impact of the coronavirus disease (COVID-19), caused by the severe acute respiratory syndrome coronavirus 2 (SARS-CoV-2), has been significant, affecting the lives of people worldwide. As of August 14th, the number of infections from this virus alone has reached approximately 595 million, with around 6 million deaths reported across 228 countries and territories ^3^. In the United States, the number of confirmed cases stands at 92,560,911, with approximately 1,031,426 lives lost as of August 2022 ^4^. The rapid spread of the coronavirus has led to the collection of a massive amount of biological data, specifically SARS-CoV-2 genomic sequencing data. This data, freely available to researchers, presents an invaluable resource for studying the virus’s behavior, developing effective vaccines, implementing preventive measures to curb transmission, and predicting the likelihood of future pandemics.

In biology, it is widely recognized that a significant portion of the mutations associated with SARS-CoV-2 primarily occurs within the spike region, also known as the peplomer protein, of the complete genome [14,3]. The structure of the SARS-CoV-2 genome, including the spike region, is depicted in Figure 1. With a genome length of approximately 30 kilobases (kb), the spike region spans the range of 21–25 kb and encodes a peplomer protein consisting of 1273 amino acids, which can be further divided into sub-units S1 and S2. Databases such as GISAID ^5^ provide freely accessible sequence data related to coronaviruses. Due to the disproportionate occurrence of mutations in the peplomer protein, analyzing protein data to gain insights into the virus’s behavior poses considerable challenges. Focusing on the peplomer protein instead of the full-length genome can save computational time when analyzing coronavirus data due to the high occurrence of mutations in this region, the sheer volume of sequences, which amounts to millions, makes it challenging to apply traditional methods like phylogenetic trees for sequence analysis [10]. Consequently, ML techniques have emerged as an appealing alternative [14,18]. However, since most ML models operate on fixed-length numerical vectors (referred to as embeddings), utilizing raw peplomer sequences as input is not feasible. Hence, the development of efficient embedding generation methods is a crucial step in ML-based classification pipelines [12]. Notable approaches, such as Spike2Vec [2] and PWM2Vec [1], have been proposed by researchers to address this challenge. Sequence alignment also plays a significant role in sequence analysis due to the importance of amino acid order. While alignment-based methods like one-hot encoding [14] have proven efficient in terms of predictive performance, there is growing interested among researchers in exploring alignment-free methods, such as Spike2Vec, to avoid the computationally expensive sequence alignment operations typically performed as a preprocessing step [5]. Alignment-free methods often employ the use of *k*-mers (or n-grams in the field of natural language processing) to generate spectra (vectors based on the frequency count of *k*-mers) [3]. Although existing alignment-free embedding methods show promising predictive results, they still pose computational challenges in terms of generation time and the high dimensionality of vectors, particularly for very long sequences and large values of *k*.

**Fig. 1:**
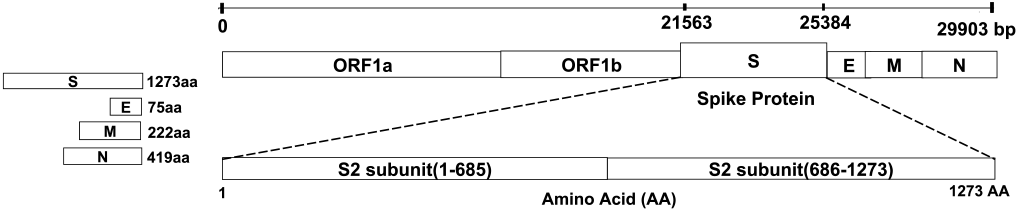
An example of SARS-CoV-2 genome. Coronavirus exhibits a dispropor-tionately high mutation rate in the S region.

In contrast to traditional embedding generation methods, which are often referred to as “feature engineering” based methods, deep learning (DL) models offer an alternative approach for sequence classification [7]. However, DL methods have not achieved significant success when it comes to classifying tabular datasets. Tree-based models like random forests and XGBoost have consistently outperformed DL methods for tabular data [4].

The use of a kernel (gram) matrix in sequence classification, especially with kernel-based machine learning classifiers like SVM, has shown promising results, as reported in prior research [8]. Kernel-based techniques have displayed favorable outcomes when compared to traditional feature engineering-based approaches [3]. These methods operate by computing kernel (similarity) values between sequences, creating a matrix based on the matches and mismatches among *k*-mers [16]. This resulting kernel matrix can also be employed for data classification with non-kernel-based classifiers, such as decision trees (by using kernel PCA [11]). However, there are two challenges associated with the kernel-based approach.

– Computing pairwise sequence similarity is expensive
– Storing *n* × *n* dimensional kernel matrix in memory (where *n* represents number of sequences) is difficult when *n* is very large. Hence, the kernel-based method cannot be scaled on a large number of sequences.

The use of the “kernel trick” can address the challenge of computing pairwise sequence similarity. However, the storage problem of storing an *n* × *n* dimensional matrix in memory remains a significant concern. To tackle these issues, our paper proposes a novel hashing-based embedding technique. This method combines the advantages of kernel methods, particularly in computing pairwise similarity between sequences, while also addressing the storage limitation. When given a peplomer protein sequence as input, our method generates embeddings by utilizing *k*-mers/nGrams (substrings of consecutive characters) and computing sequence sketches, followed by a dot product operation to avoid full spectrum computation (frequency count). By operating on low-dimensional vectors (sketches) instead of an *n* × *n* matrix or full-length spectrum encompassing all possible *k*-mer combinations, our method enables scalability to handle a large number of sequences, effectively resolving the scalability problem. Our fast and alignment-free method seamlessly integrates with machine learning algorithms. It supports classification and clustering tasks, including k-nearest neighbors and decision trees. This paper presents the following contributions:

1. We introduce a fast, alignment-free, and efficient embedding method that rapidly computes low-dimensional numerical embeddings for protein sequences.
2. Our method combines kernel method benefits, addressing the scalability challenge for pairwise similarity.
3. Our Experiment achieves up to 99.6% faster embedding generation reduction in computation time compared to state-of-the-art (SOTA) methods.
4. Our results show that our method outperforms existing methods, achieving accuracy and ROC-AUC scores of up to 86% and 85%, respectively.
5. Visualization shows the similarity of our method’s embeddings to SOTA.

In the upcoming sections, we provide literature review in Section 2, proposed model ins Section 3, experimental setup in Section 4, results in Section 5, and conclusion in Section 6.

## 2 Related Work

The feature engineering-based methods, such as Spike2Vec [2] and PWM2Vec [1], leverage the concept of using *k*-mers to achieve satisfactory predictive performance. However, these methods encounter the challenge known as the “curse of dimensionality.” Specifically, as the value of *k* increases, the resulting spectrum (frequency count vector) becomes increasingly sparse, with smaller *k*-mers occurring less frequently. Consequently, the likelihood of encountering a specific *k*-mer again decreases. To address this issue of sparse vectors, the authors in [9] propose the utilization of gapped or spaced *k*-mers. Gapped *k*-mers involve extracting *g*-mers from larger *k*-mers, where *g* is a value smaller than *k*. In this approach, a *g*-mer consists of the first *g* characters (amino acids), while the remaining characters are disregarded.

The computation of pair-wise similarity between sequences using kernel matrices has been extensively studied in the field of ML [13], which can be computationally expensive. To address this issue, an approximate method was proposed by authors in [8], involving computing the dot product between the spectra of two sequences. The resulting kernel matrix can then be utilized as input for SVM or non-kernel classifiers using kernel PCA [11] for classification purposes. In a different approach, authors in [19] introduced a neural network-based model that employs the Wasserstein distance (WD) to extract features. It is worth noting that feature embeddings have applications in various domains, including product recommendations [12].

Feature engineering-based methods, while achieving improved predictive performance, often struggle to generalize effectively across different types of input sequences. Deep learning-based methods offer a potential solution to this problem [4]. In [22], authors use the ResNet for classification. However, deep learning methods generally do not yield promising results when applied to tabular data [20]. Converting protein sequences into images for input in existing image classification deep learning models is an alternative approach [17], yet preserving all pertinent information during the transformation remains a challenging task.

## 3 Proposed Approach

In this section, we delve into the specifics of our sequence representation and conduct an analysis of the computational complexities involved. As mentioned earlier, sequences typically exhibit varying lengths, and even when their lengths are identical, they might not be aligned. Consequently, treating them as straight-forward vectors is not feasible. Even in the case of aligned sequences with equal lengths, treating them as vectors fails to account for the sequential order of elements and their continuity. To address these challenges comprehensively, one of the most successful approaches involves representing sequences with fixed-dimensional feature vectors. These feature vectors comprise the spectra or counts of all *k*-mers, which are strings of length *k*, found within the sequences [15].

Consider a scenario where we have at our disposal a collection of spike/peplomer protein sequences denoted as *𝒮*, composed of amino acids. Now, for a fixed positive integer *k*, let’s define *Σ*^*k*^ as the set comprising all strings with a length of *k* formed from characters in *Σ* (essentially, all conceivable *k*-mers). Consequently, there would be a total of *s* = |*Σ*|^*k*^ possible *k*-mers in this set. The spectrum, denoted as *Φ*_*k*_(*X*), associated with a sequence *X* ∈ *𝒮* can be envisioned as a vector spanning *s* dimensions. Each dimension corresponds to the count of a specific *k*-mer occurring within the sequence *X*. In more precise terms,

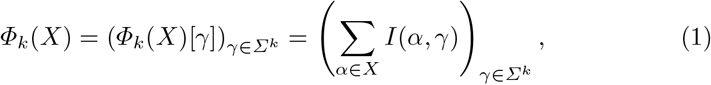

Where

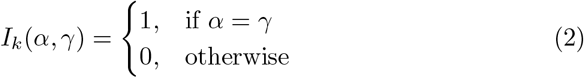

It’s essential to note that *Φ*_*k*_(*X*) represents a vector with *s* = |*Σ*|^*k*^ dimensions. In this vector, each coordinate, denoted as *γ* and belonging to the set *Σ*^*k*^, holds a value equal to the frequency of *γ* within the sequence *X*. Specifically, for peplomer sequences, where |*Σ*| = 20, the length of the feature vector grows exponentially with increasing *k*. However, in the case of other sequences like discretized music signals or text, *Σ* might be considerably larger, leading to a substantial increase in the space required to represent these sequences, which can become impractical.

In the realm of kernel-based machine learning, a vital component is the “kernel function.” This function calculates a real-valued similarity score for a pair of feature vectors. Typically, this kernel function computes the inner product of the respective spectra.

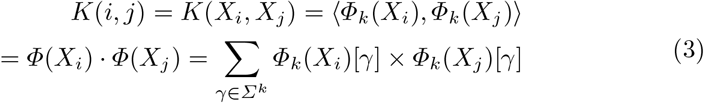

The utilization of a kernel matrix, also known as pairwise similarities, serves as input for a conventional support vector machine (SVM) classifier. This approach has proven to deliver outstanding classification performance across various applications [8]. However, the so-called *kernel trick* aims to bypass the explicit computation of feature vectors. While this technique is advantageous in terms of avoiding the quadratic space requirements for storing the kernel matrix, it encounters scalability issues when dealing with real-world sequence datasets.

### Proposed Representation

In our proposed approach, denoted as 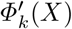 for a sequence *X*, we provide an approximation of the feature vector *Φ*_*k*_(*X*). This approximation enables the application of machine learning methods based on vector space. Importantly, we establish a close relationship between the Euclidean distance of a pair of vectors and the aforementioned kernel-based proximity measure. For a sequence *X* within the set *𝒮*, we represent 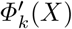 as an approximate form of the spectrum *Φ*_*k*_(*X*). To calculate *Φ*′(·), we employ 2-universal hash functions.

#### Definition 1 (2-universal hash function)

*A family* ℌ *of functions of the form h* : *Σ*^*k*^ ↦ [*w*] *is called a* 2*-universal family of hash functions, if for a randomly chosen h* ∈ ℌ

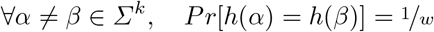

#### Definition 2 (Linear Congruential hash functions)

*For an integer w, let p > w be a large prime number. For integers* 0 *< a < p and* 0 ≤ *b < p, and α* ∈ *Σ*^*k*^ *(represented as integer base* |*Σ*|*), the hash function h*_*a,b*_ : *Σ*^*k*^ *↦* [*w*] *is defined as*

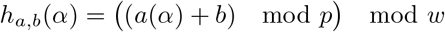

It is well-known that the family ℌ = {*h*_*a,b*_ : 0 *< a < p*, 0 ≤ *b < p*} is 2-universal. For an 0 *< ϵ <* 1, let *w >* 2*/ϵ* be an integer. Suppose 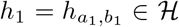 is randomly selected. 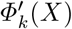 is a *w*-dimensional vector of integers. The *i*th coordinate of 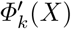 is the cumulative frequency of all *k*-mers *α* that hash to the bucket *i* by *h*_1_, i.e.

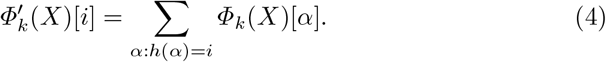

Next, we show that the dot-product between the approximate representation of a pair of sequences *X* and *Y* closely approximates the kernel similarity value given in (3). Then we extend this basic representation using multiple hash functions to amplify the goodness of the estimate.

We are going to show that for any pair of sequences *X* and *Y*, 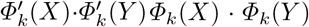. For notational convenience let 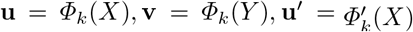, and 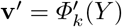, we show that **u**′ · **v**′ ≃ **u ·v**.

#### Theorem 1 u ·v ≤ u′ · v′ ≤ **u ·v** + *ϵ*∥u∥_1_∥v∥_1_ *with probability* ≥ ½

***Theorem 1 Proof:***

*Proof*.

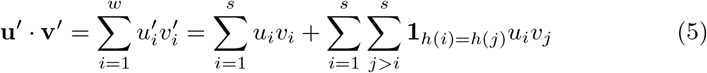

where **1**_*h*(*i*)=*h*(*j*)_ is the indicator function for the event *h*(*i*) = *h*(*j*). Since entries in **u** and **v** are non-negative integers, we get that the first inequality holds certainly. For the second inequality, we estimate the error term 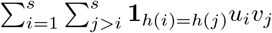

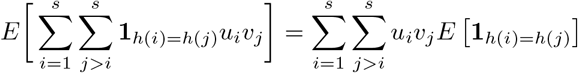

By the 2-university of *h*, we get *E*[**1**_*h*(*i*)=*h*(*j*)_] = 1*/w* Using *w* ≥ 2*/ϵ*, we get that

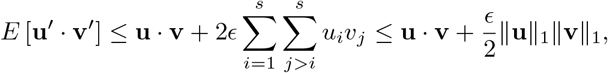

where the last inequality uses the Cauchy-Shwarz inequality. By the Markov inequality, with probability at most 1*/*2 the error is more than *ϵ*∥**u**∥_1_ ∥**v**∥_1_, hence the statement of the theorem follows. □

### Mathematical Proofs

1. asserts that 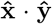 is an unbiased estimate for the kernel similarity. While 2. provides a bound on the deviation of the estimate.

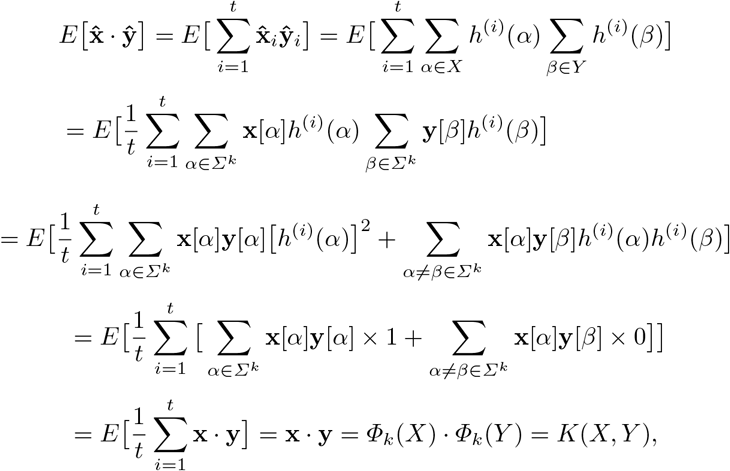

Note that the upper bound on the error is very loose, in practice we get far better estimates of the inner product. In order to enhance the result (so the error is concentrated around its mean), we use *t* hash functions *h*_1_, …, *h*_*t*_ from the family *ℌ*. This amounts to randomly choosing *t* pairs of the integers *a* and *b* in the above definition. In this case, our representation for a sequence *X* is the scaled concatenation of *t* elementary representations. For 1 ≤ *i* ≤ *t*, let 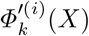 be the representation under hash function *h*_*i*_ (from Equation (4)). Then,

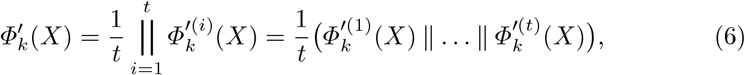

where ‖ is the concatenation operator. The quality bound on the approximation of (3) holds as in Theorem 1. Note that the definition of our representation of (4) is derived from the count-min sketch of [6], except for they take the minimum of the inner products over the hash functions and attain a better quality guarantee. Since we want a vector representation, we cannot compute the *“non-linear”* functions of min and median.

#### Remark 1

It’s worth highlighting that our representation as expressed in (6) for a given sequence *X* can be computed efficiently in a single linear scan over the sequence, as detailed in Algorithm 1. Consequently, the runtime required to compute 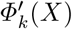 is proportional to *tn*_*x*_, where *n*_*x*_ denotes the number of characters in sequence *X*. Regarding the space complexity associated with storing 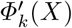, it can be described as 2*t/ϵ*, with both *ϵ* and *t* being parameters under the user’s control. Additionally, it’s important to note that within the error term, |**u**|_1_ = *n*_*x*_ − *k* + 1, with **u** representing the spectrum of the sequence *X*.

Subsequently, we demonstrate a close connection between the Euclidean or *ℓ*_2_-distance commonly utilized in vector-space machine learning techniques such as *k*-NN classification or *k*-means clustering and the kernel similarity score defined in (3). This alignment allows our approach to harness the advantages of kernel-based learning without incurring the time complexity associated with kernel computation and the space complexity required for storing the kernel matrix. Through appropriate scaling of the vectors 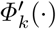, we derive detailed evaluations, which can be found in the supplementary material.

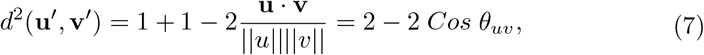

where **u, u**′ are convenient notation for *Φ*_*k*_(*X*) and 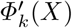, as defined above and *d*(·,·) is the Euclidean distance. Thus, both the “Euclidean and cosine similarity” are proportional to the ‘kernel similarity’.

The pseudocode of our method is given in Algorithm 1. For simplicity, we use a Python-like style for the pseudocode. Our algorithm takes a set of peplomer sequences *S*, integer *k, m, p*, alphabet *Σ*, and number of hash functions *h*. It outputs a sketch *Φ*, which is a low dimensional fixed-length numerical embedding corresponding to each peplomer sequence. The method starts by initializing *Φ* as an empty array (line 4), *m* with 2^10^ in line 5 (where we can take any integer power of 2), and *p* with 4999 (where *p* is any four-digit prime number and *p > m*). Now we iterate each sequence one by one (line 7) and generate a set of *k*-mers (line 8). Note that we take *k* = 3 here using the standard validation set approach. In the next step, for multiple hash functions *h*, where *h* is any integer value ≥ 1, we need to compute a numerical representation for each *k*-mer and store their frequencies in a local sketch list. The length of the local sketch list (for each sequence) equals to *m* (line 12). Since our idea to store the values in sketch is based on hashing, we initialize two variables *a*1 (random integer between 2 and *m* − 1) and *b*1 (random integer between 0 and *m* − 1) in line 13 and 14. To compute the integer number corresponding to each *k*-mer, we first compute each *k*-mers characters (amino acids) positions in the alphabet *Σ* (line 18), where *Σ* comprised of 21 characters *ACDEFGHIKLMNPQRSTVWXY*. We then sort the characters in *k*-mers (line 19) and note their position (line 20). Finally, we assign a numerical value to a character, which comprised of its position in *Σ* times |*Σ*| ^its position in the *k*-mer^ (line 21). This process is repeated for all characters within a *k*-mer (loop in line 17) and storing a running sum to get an index value for any given *k*-mer. Similarly, the same process is repeated for all *k*-mers within a sequence (loop in line 15). Now, we define the hash function (line 23):

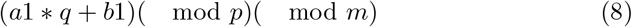

where *q* is the integer value assigned to the *k*-mer. After getting the hash value using Equation 8, we increment that index (integer hash value) in the local sketch array by one (line 24). This process is repeated for all *k*-mers within a sequence. We normalize the local sketch list by first dividing it by the total sum of all values in the list and then dividing it by *h*, which is the number of hash functions (line 26). This process is repeated *h* times (loop in line 11) for all hash functions and all local sketch lists are concatenated to get the final sketch *Φ*_*s*_ for a single sequence.

## 4 Experimental Evaluation

In this section, we discuss the dataset used for experimentation and introduce state-of-the-art (SOTA) methods for comparing results. In the end, we show the visual representation of the proposed SOTA embeddings to get a better understanding of the data. All experiments are performed on a core i5 system (with a 2.40 GHz processor) having Windows OS and 32 GB memory. For experiments, we use the standard 70-30% split for training and testing sets, respectively. We repeat each experiment 5 times to avoid randomness and report average results.

### Dataset Statistic

We extracted the spike sequences from a popular database called GISAID ^6^. The total number of sequences we extracted is 7000, representing 22 coronavirus variants (class labels) [3].

#### Algorithm 1 Proposed method computation

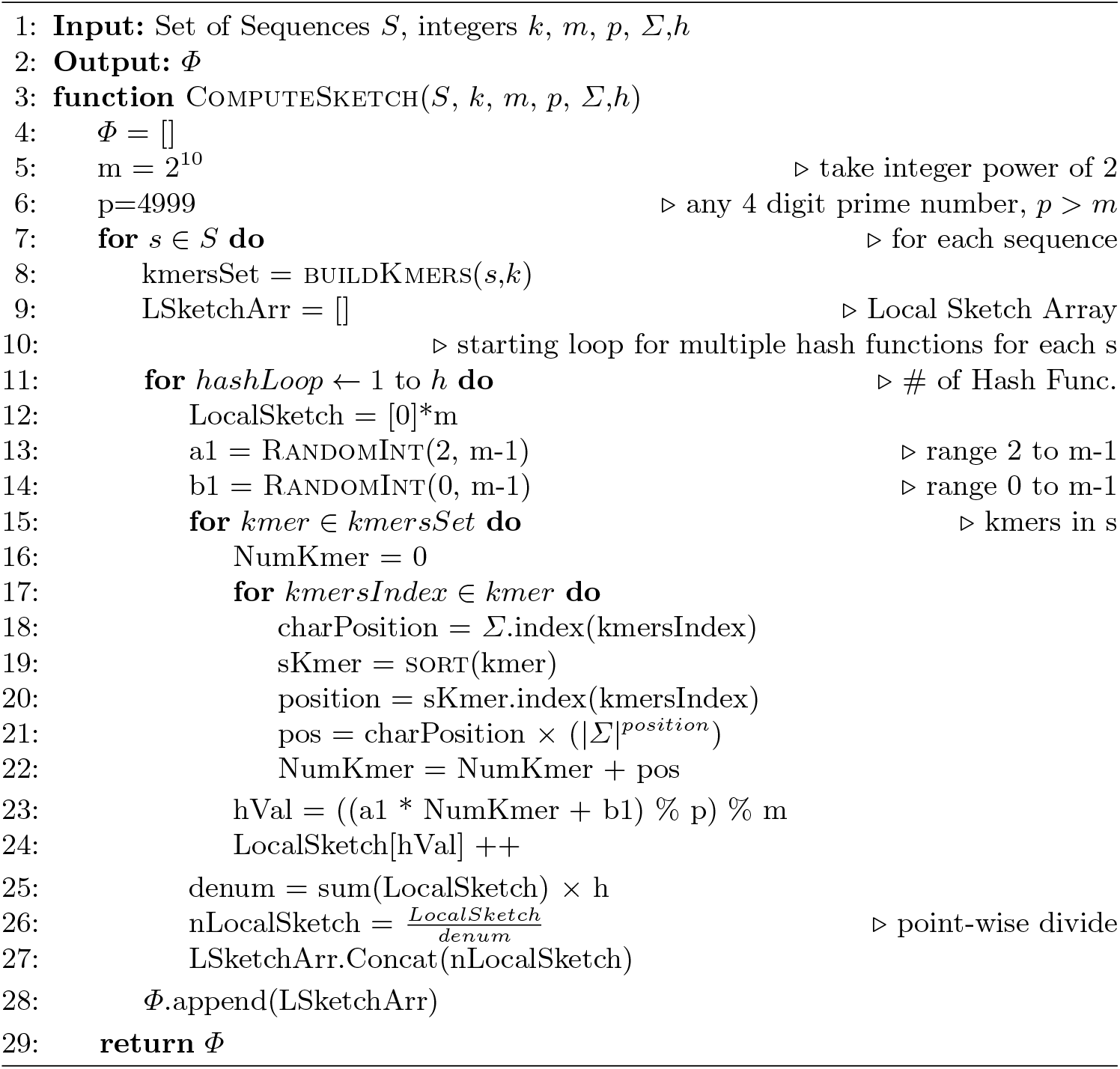

#### Remark 2

Note that we use the lineages (i.e., B.1.1.7, B.1.617.2, etc., as the class labels for classification purposes). That is, given the embeddings as input, the goal is to classify the lineages using different ML classifiers.

For classification purposes, our study employs a variety of classifiers, including Support Vector Machine (SVM), Naive Bayes (NB), Multi-Layer Perceptron (MLP), K-Nearest Neighbors (KNN), Random Forest (RF), Logistic Regression (LR), and Decision Tree (DT) models. To assess the performance of these diverse models, we employ a range of evaluation metrics, including average accuracy, precision, recall, weighted F1 score, macro F1 score, Receiver Operator Characteristic Curve (ROC) Area Under the Curve (AUC), and training runtime. In scenarios where metrics are designed for binary classification, we employ the one-vs-rest strategy for multi-class classification.

### State-of-the-Art (SOTA) Models

We use five state-of-the-art methods, namely Spike2Vec [2], PWM2Vec [1], String Kernel [8], Wasserstein Distance Guided Representation Learning (WDGRL) [19], and Spaced *k*-mers [21], for comparison of results.

## 5 Results and Discussion

In this section, we report the classification results for our method and compare the results with SOTA methods.

### Increasing Number of Hash Functions

Results for our method with an increasing number of hash functions is shown in Table 1. In this experimental setting, we use *k* = 3 for the *k*-mers. We can observe that although there is not any drastic change in results among different numbers of hash functions *h*, the random forest classifier with *h* = 2 outperforms all other classifiers and values of *h* for all but one evaluation metric. For training time, since the sketch length for *h* = 1 is the smallest among the others, it took the least amount of time to train classifiers.

**Table 1:** Classification results showing the effect of changing number of hash functions with k=3 for *k*-mers. Best values are shown in bold.

| Parameter<br>h | Algo. | Acc. | Prec. | Recall | F1<br>(Weig.) | F1<br>(Macro) | ROC<br>AUC | Train Time<br>(sec.) |
| --- | --- | --- | --- | --- | --- | --- | --- | --- |
| Number<br>of Hash<br>Func-<br>tions: 1 | SVM | 0.845 | 0.848 | 0.845 | 0.836 | 0.675 | 0.837 | 4.034 |
|  | NB | 0.612 | 0.739 | 0.612 | 0.635 | 0.447 | 0.731 | **0.708** |
|  | MLP | 0.819 | 0.820 | 0.819 | 0.813 | 0.604 | 0.800 | 12.754 |
|  | KNN | 0.806 | 0.821 | 0.806 | 0.801 | 0.616 | 0.797 | 0.965 |
|  | RF | 0.854 | 0.855 | 0.854 | 0.844 | 0.680 | 0.836 | 1.705 |
|  | LR | 0.482 | 0.243 | 0.482 | 0.318 | 0.030 | 0.500 | 3.492 |
|  | DT | 0.841 | 0.844 | 0.841 | 0.833 | 0.663 | 0.829 | 0.327 |
| Number<br>of Hash<br>Func-<br>tions: 2 | SVM | 0.848 | 0.858 | 0.848 | 0.841 | 0.681 | 0.848 | 9.801 |
|  | NB | 0.732 | 0.776 | 0.732 | 0.741 | 0.555 | 0.771 | 1.440 |
|  | MLP | 0.835 | 0.825 | 0.835 | 0.825 | 0.622 | 0.819 | 13.893 |
|  | KNN | 0.821 | 0.818 | 0.821 | 0.811 | 0.616 | 0.803 | 1.472 |
|  | RF | **0.863** | **0.867** | **0.863** | **0.854** | **0.703** | **0.851** | 2.627 |
|  | LR | 0.500 | 0.264 | 0.500 | 0.333 | 0.031 | 0.500 | 11.907 |
|  | DT | 0.845 | 0.856 | 0.845 | 0.841 | 0.683 | 0.839 | 0.956 |
| Number<br>of Hash<br>Func-<br>tions: 3 | SVM | 0.842 | 0.845 | 0.842 | 0.832 | 0.678 | 0.840 | 14.189 |
|  | NB | 0.639 | 0.741 | 0.639 | 0.655 | 0.474 | 0.736 | 2.100 |
|  | MLP | 0.817 | 0.816 | 0.817 | 0.809 | 0.608 | 0.802 | 18.490 |
|  | KNN | 0.811 | 0.812 | 0.811 | 0.804 | 0.616 | 0.797 | 1.981 |
|  | RF | 0.852 | 0.852 | 0.852 | 0.841 | 0.689 | 0.842 | 2.966 |
|  | LR | 0.482 | 0.233 | 0.482 | 0.314 | 0.030 | 0.500 | 8.324 |
|  | DT | 0.841 | 0.846 | 0.841 | 0.833 | 0.679 | 0.837 | 1.279 |

### Comparison With SOTA

A comparison of our method with SOTA algorithms is shown in Table 2. For these results, we report our method results for *h* = 2 because it showed best performance in Table 1. We can observe that the proposed method (with *h* = 2 and *k* = 3) outperforms all the SOTA methods for all but one evaluation metric. In the case of training runtime, WDGRL performs the best.

**Table 2:**
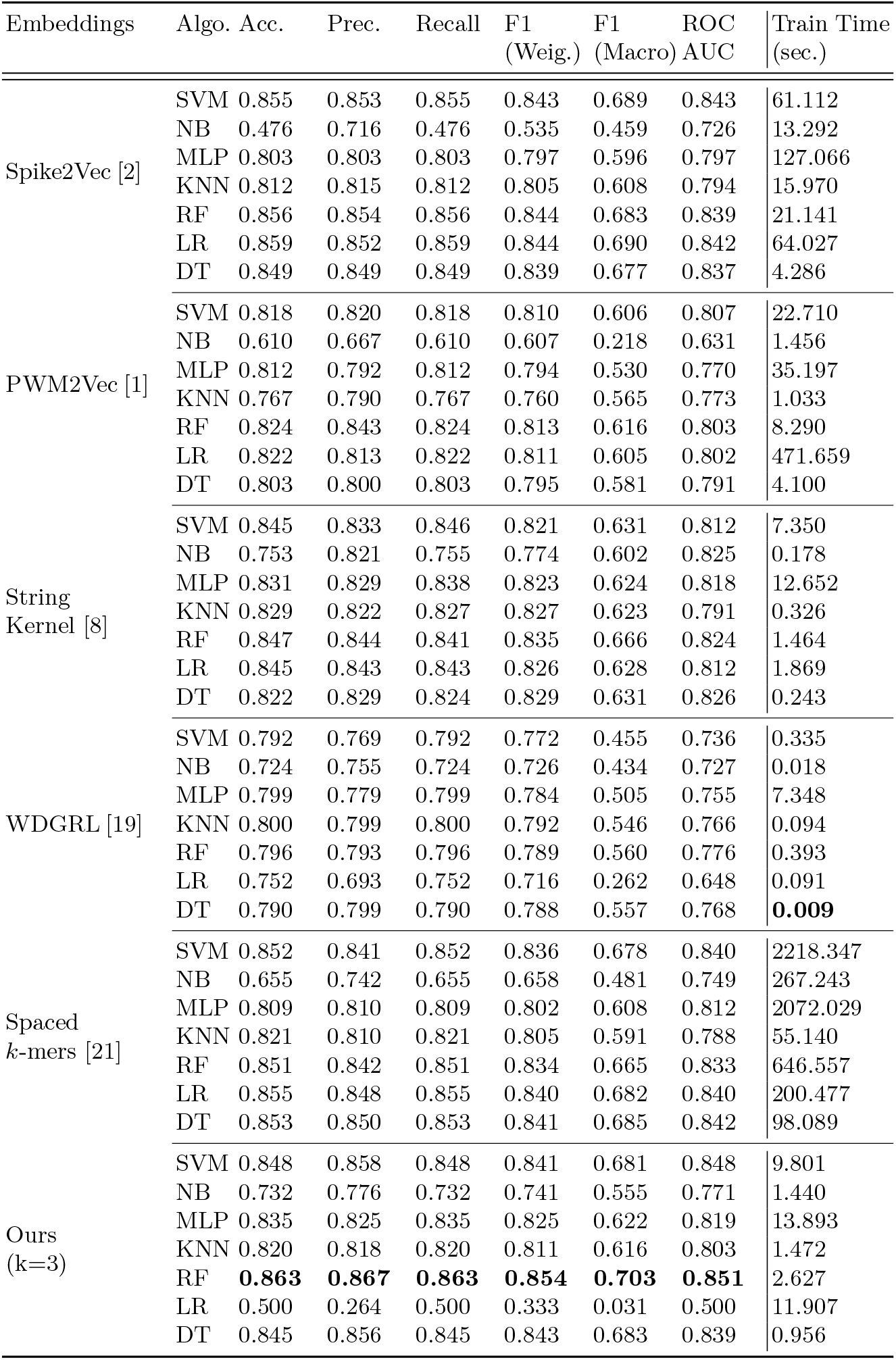
Classification performance of SOTA and Our methods.

### Effect of k for *k*-mers

To evaluate the effect of *k* for *k*-mers on the results of our method, we report the results of our model with varying values of k in Table 3. We can observe that RF classifier with *k* = 3 outperforms other values of *k* for all but one evaluation metric. In terms of training runtime, decision tree classifiers take the least time.

**Table 3:** Classification results showing the effect of k for *k*-mers with *h* = 2 for proposed method. The best values are shown in bold.

| Parameter<br>k | Algo. | Acc. | Prec. | Recall | F1<br>(Weig.) | F1<br>(Macro) | ROC<br>AUC | Train Time<br>(sec.) |
| --- | --- | --- | --- | --- | --- | --- | --- | --- |
| k=3 | SVM | 0.848 | 0.858 | 0.848 | 0.841 | 0.681 | 0.848 | 9.801 |
|  | NB | 0.732 | 0.776 | 0.732 | 0.741 | 0.555 | 0.771 | 1.440 |
|  | MLP | 0.835 | 0.825 | 0.835 | 0.825 | 0.622 | 0.819 | 13.893 |
|  | KNN | 0.821 | 0.818 | 0.821 | 0.811 | 0.616 | 0.803 | 1.472 |
|  | RF | **0.863** | **0.867** | **0.863** | **0.854** | **0.703** | **0.851** | 2.627 |
|  | LR | 0.500 | 0.264 | 0.500 | 0.333 | 0.031 | 0.500 | 11.907 |
|  | DT | 0.845 | 0.856 | 0.845 | 0.841 | 0.683 | 0.839 | **0.956** |
| k=5 | SVM | 0.850 | 0.847 | 0.850 | 0.836 | 0.680 | 0.839 | 8.827 |
|  | NB | 0.640 | 0.715 | 0.640 | 0.640 | 0.463 | 0.721 | 1.432 |
|  | MLP | 0.826 | 0.823 | 0.826 | 0.816 | 0.629 | 0.813 | 13.375 |
|  | KNN | 0.818 | 0.824 | 0.818 | 0.812 | 0.621 | 0.801 | 1.319 |
|  | RF | 0.857 | 0.853 | 0.857 | 0.843 | 0.690 | 0.842 | 2.322 |
|  | LR | 0.483 | 0.237 | 0.483 | 0.315 | 0.030 | 0.500 | 7.219 |
|  | DT | 0.844 | 0.840 | 0.844 | 0.833 | 0.667 | 0.834 | 0.987 |
| k=7 | SVM | 0.853 | 0.854 | 0.853 | 0.841 | 0.691 | 0.846 | 9.782 |
|  | NB | 0.642 | 0.721 | 0.642 | 0.644 | 0.452 | 0.721 | 1.398 |
|  | MLP | 0.831 | 0.826 | 0.831 | 0.821 | 0.634 | 0.818 | 13.363 |
|  | KNN | 0.823 | 0.827 | 0.823 | 0.817 | 0.637 | 0.816 | 1.378 |
|  | RF | 0.856 | 0.854 | 0.856 | 0.844 | 0.692 | 0.845 | 2.644 |
|  | LR | 0.485 | 0.236 | 0.485 | 0.317 | 0.030 | 0.500 | 7.942 |
|  | DT | 0.842 | 0.841 | 0.842 | 0.833 | 0.656 | 0.830 | 1.090 |
| k=9 | SVM | 0.849 | 0.847 | 0.849 | 0.838 | 0.676 | 0.836 | 10.099 |
|  | NB | 0.644 | 0.714 | 0.644 | 0.651 | 0.437 | 0.707 | 1.540 |
|  | MLP | 0.833 | 0.830 | 0.833 | 0.825 | 0.625 | 0.810 | 12.938 |
|  | KNN | 0.820 | 0.826 | 0.820 | 0.815 | 0.622 | 0.802 | 2.842 |
|  | RF | 0.853 | 0.852 | 0.853 | 0.842 | 0.679 | 0.835 | 3.127 |
|  | LR | 0.485 | 0.236 | 0.485 | 0.317 | 0.030 | 0.500 | 8.140 |
|  | DT | 0.836 | 0.836 | 0.836 | 0.828 | 0.647 | 0.821 | 1.127 |

### Embeddings Generation Time

To illustrate the efficiency of our approach regarding embedding computation time, we conducted a runtime comparison with state-of-the-art (SOTA) methods, as summarized in Table 4. Our proposed method stands out by requiring the shortest time, completing the feature vector (sketch) generation in just 47.401 seconds, outperforming the SOTA alternatives. Among the alternatives, PWM2Vec, while not an alignment-free approach, emerges as the second-best in terms of runtime. In contrast, generating feature vectors with spaced *k*-mers consumes the most time. We also present the percentage improvement in runtime achieved by our method compared to PWM2Vec (the second-best in runtime) and Spaced *k*-mers (the slowest). To calculate this improvement, we use the formula: 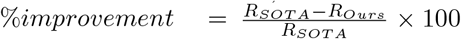. Here, *R*_SOTA_ represents the runtime of SOTA embedding methods (PWM2Vec and Spaced *k*-mers), while *R*_Ours_ corresponds to the runtime of our method’s embedding computation. Table 4 clearly illustrates that our proposed method enhances runtime performance significantly, improving it by 70.9% and 99.6% compared to PWM2Vec and Spaced *k*-mers, respectively.

**Table 4:** Embedding generation runtime for different methods. The best value is shown in bold. The percentage improvement of the runtime is also given for our method.

| Embeddings | Runtime (Sec.) |
| --- | --- |
| Spike2Vec [2] | 354.061 |
| PWM2Vec [1] | 163.257 |
| String Approx. [8] | 2292.245 |
| WDGRL [19] | 438.188 |
| Spaced $k$-mers [21] | 12901.808 |
| **Ours** | **47.401** |
| % Improv. of our method from PWM2Vec | 70.9% |
| % Improv. of our method from Spaced $k$-mers | 99.6% |

## 6 Conclusion

This paper introduces a novel approach for rapidly generating protein sequence sketches that are both efficient and alignment-free, leveraging the concept of hashing. Our method not only exhibits swift sketch generation but also enhances classification outcomes when compared to existing methodologies. To validate our model, we conducted extensive experiments using real-world biological protein sequence datasets, employing a variety of evaluation metrics. Our method demonstrates an impressive 99.6% enhancement in embedding generation runtime compared to the state-of-the-art (SOTA) approach. Future endeavors will involve assessing our method’s performance on more extensive sets of sequence data, potentially reaching multi-million sequences. Additionally, we aim to apply our approach to other virus data, such as Zika, to further explore its utility and effectiveness.

## Footnotes

3 https://www.worldometers.info/coronavirus/

4 https://www.cdc.gov/coronavirus/2019-ncov/index.html

5 https://gisaid.org/

6 https://www.gisaid.org/

